# Design, Performance Evaluation and Investigation of the Dynamic Mechanisms of Earthworm-Microorganisms interactions for wastewater treatment through Vermifiltration technology

**DOI:** 10.1101/2020.08.14.252072

**Authors:** Sudipti Arora, Sakshi Saraswat, Rinki Mishra, Jayana Rajvanshi, Jasmine Sethi, Aditi Nag, Sonika Saxena

## Abstract

The present study points to the relevance of the earthworms-microorganism’s symbiotic and synergistic interactions that drive the wastewater treatment by identifying the most essential mechanisms underlying the removal of contaminants during vermifiltration technology. Previous studies have showed the presence of earthworms improves treatment performance of vermifilter (VF), but earthworm microbial community dynamics, their structure and functional characteristics in VF were not fully investigated. To investigate the effects of earthworms on the inherent microbial community of the VF, the present study envisages the dynamics of the complex symbiosis of earthworms & microorganisms associated to the treatment mechanisms. In this study, the design, operations and performance evaluation and influent, effluent and filter media layer were investigated for microbial diversity inside the earthworm population, along with the antimicrobial activity, enzymatic activity, and protein profiling assays. The results showed that earthworm gut microbial communities were dominated by *Gammaproteobacteria*, and the percentages arrived to 59–60% of the microbial species detected, while filter media layer showed presence of *Firmicutes* and *Actinobacteria*. The protein profiling of the microbiota associated with the VF showed that earthworms feeding and earthworm–microorganism interaction were responsible for enhanced treatment performance. The finding provides an insight into the complex earthworm microbial dynamics and mechanisms for wastewater treatment in VF. Furthermore, earthworm predation strongly regulated microbial biomass while improving microbial activity, and is deciphered as the possible mechanisms behind the vermifiltration technology.

## 1. Introduction

In a developing country like India, the uncontrolled growth in urban areas has left many cities deficient in water supply, sewerage, and storm water drainage services and it is due to these deficient services that sewage and its management has become a tenacious problem, even though a large part of the municipal expenditure is allotted to it (Kumar and Joseph, 2012). This, in turn, results in an increase in morbidity especially due to pathogens and parasitic infections and infestations in all segments of the population, particularly the urban slum dwellers. The main cause of water pollution is the unintended disposal of untreated, partly treated and non-point sources of sewage and more important is its effect on human health and environment. It has also been observed that some sewage treatment plants (STPs) do not meet prescribed standards concerning Biochemical Oxygen Demand (BOD) thereby rendering the treated water unsuitable for reuse purpose along with the huge operation and maintenance expenditures. Thus, instead of traditional setting up of STPs, it is important to now look at alternative methods of wastewater recycling *viz*. decentralized, sustainable and economical technologies which do not burden the economy of developing countries. Considering the current challenges in access and management of conventional sanitation, it is necessary to explore low-cost technology and systems that provide a closed-loop between health and financial system, promoting circular economy. Such technology must include simple operation and maintenance, high treatment efficiency, ecological, innovative, sustainable option along with the potential for sub-products reuse (Li et al., 2009) and full integration of faecal sludge (FS) management.

Worldwide, the potential of using earthworms (EWs) to treat municipal sewage sludge, domestic wastewater, and human faeces is increasing, and many previous research studies have already shown that vermifiltration-based on-site sanitation could constitute alternatives to existing municipal and domestic wastewater treatment as well as faecal waste treatment technology (Zhao et al., 2010; Furlong et al., 2015). Vermifiltration is nature-based sanitation solution for wastewater treatment technology, where polluted wastewater is treated with the help of earthworms resulting in treated effluent and organic manure or vermicompost as useful byproducts. Vermifiltration is a bio-oxidative process in which earthworms interact intensively with microorganisms within the decomposer community, accelerating the stabilization of organic matter and greatly modifying its physical and biochemical properties (Dominguez and Edwards, 2004). Although microorganisms are primarily responsible for the biochemical degradation of organic matter, microbial communities developed in the VF due to the activity of earthworms in the filter bed, are responsible for different mechanisms, mainly modification of microbial biomass, as well as microbial community structure and functional activity through burrowing, digesting and dispersing microbes, and casting behavior (Brown, 1995; Aira and Domínguez, 2011). Due to their importance in soil ecosystem, earthworm casts have been widely studied in the past, when seeking to understand changes in the nutrient status and microbial composition of the soil (Furlong et al., 2002; Aira and Domínguez, 2011). It follows that earthworm casts may exert direct and indirect effects on the microbial communities of biofilms in the VF. Additionally, the microorganisms in earthworm gut play an important role in the digestion of organic food. Xing et al., 2010 revealed that passage of the intestinal tract of earthworm had a qualitative and quantitative influence on the microbial community of the VF biofilm. However, the knowledge about microbial community composition in earthworm gut is limited. There are few studies that suggest that earthworms’ body works as a ‘biofilter’ and they have been found to remove contaminants by 80-90% from wastewater (Sinha et al., 2008). It is also reported that earthworms create aerobic conditions in the waste materials by their burrowing actions, inhibiting the action of anaerobic microorganisms and the enhanced performance of the vermifiltration process for wastewater is due to better aerobic conditions from the burrowing action of earthworms, the greater adsorption effect of the earthworm casts, and higher levels of microbial activity stimulated earthworm feeding (Sinha et al., 2008; Zhao et al., 2010). There are very few studies that shed light on the probable mechanism behind the treatment of wastewater by VF. Very little attention has been paid so far to the impact of earthworms on the microbial community structure and function in VF. In particular, the influence of earthworm castings on the microbial community in VF biofilms is poorly understood. It is important to understand that microorganisms play an important role in the removal of organic load during wastewater treatment in a biofilter (which is devoid of EWs) but the major dynamics or changes after adding earthworms into the VF system which leads to enhanced treatment is still not known. Therefore, for successful implementation of VF technology, we need to decipher the complex symbiotic relationship between earthworms and microorganisms and their respective roles in wastewater treatment. The present study attempts to elucidate the above-stated mechanism by designing, evaluating the performance efficacy of VF technology for the treatment of institutional wastewater and understanding the microbial community dynamics for wastewater treatment. The objective of the study is to investigate the microbial community diversity from influent, effluent, earthworms’ whole body and casts, EW’s coelomic fluid and filter media layer (active layer) of the VF and determine their antimicrobial and enzymatic activity to decode their respective roles for the treatment. Thus, investigation of the microbial consortium and functionality in a VF can help in understanding and controlling the vermifiltration process, and resulting in the overall optimization of the vermifiltration system.

## 2. Methods

### 2.1. Vermifiltration setup & operation

A vermifilter of capacity (1 m^3^) was installed at the field site at the campus of Dr. B. Lal Institute of Biotechnology, Jaipur (India) for the treatment of institutional wastewater (as represented in Figure 1) and was allowed to operate for a period of one year. All the parameters were selected based on previous findings (Arora et al., 2016, 2014b) considering the climatic conditions of Jaipur, Rajasthan. A layer of natural fiber was placed on top of the filter bed to avoid direct hydraulic influence on the earthworms The wastewater was collected from the campus in a collection tank of capacity 500 liters and was then pumped to an overhead tank of capacity 300 liter. The wastewater was allowed to mix in the overhead tank with the help of an agitator and this wastewater was allowed to flow by sprinkler through gravity at HLR of 1.0 m^3^/m^2^/d and HRT of 4-6 hours. The active layer consisted of mature cow dung vermigratings, in which *Eisenia fetida* were inoculated with a stocking density of 10,000 worms/ m^3^. These layers were chosen based on previous studies (Arora et al., 2014b, 2014a; Arora and Kazmi, 2015), which helped in providing proper filtration, and bedding to allow earthworms to interact with microorganisms. Initially, the VF was allowed to operate for a period of 15 days to acclimatize the whole setup. Then earthworms were inoculated in the active layer and the VF was monitored for an initial acclimatization period of 21 days where the earthworms were allowed to adapt to the changing environment. Some EWs were not added to the VF and were used as control experiments for elucidating the mechanistic insights. After acclimation for 21 days, the VF was operated continuously for 55 weeks at natural field conditions.

**Figure 1:**
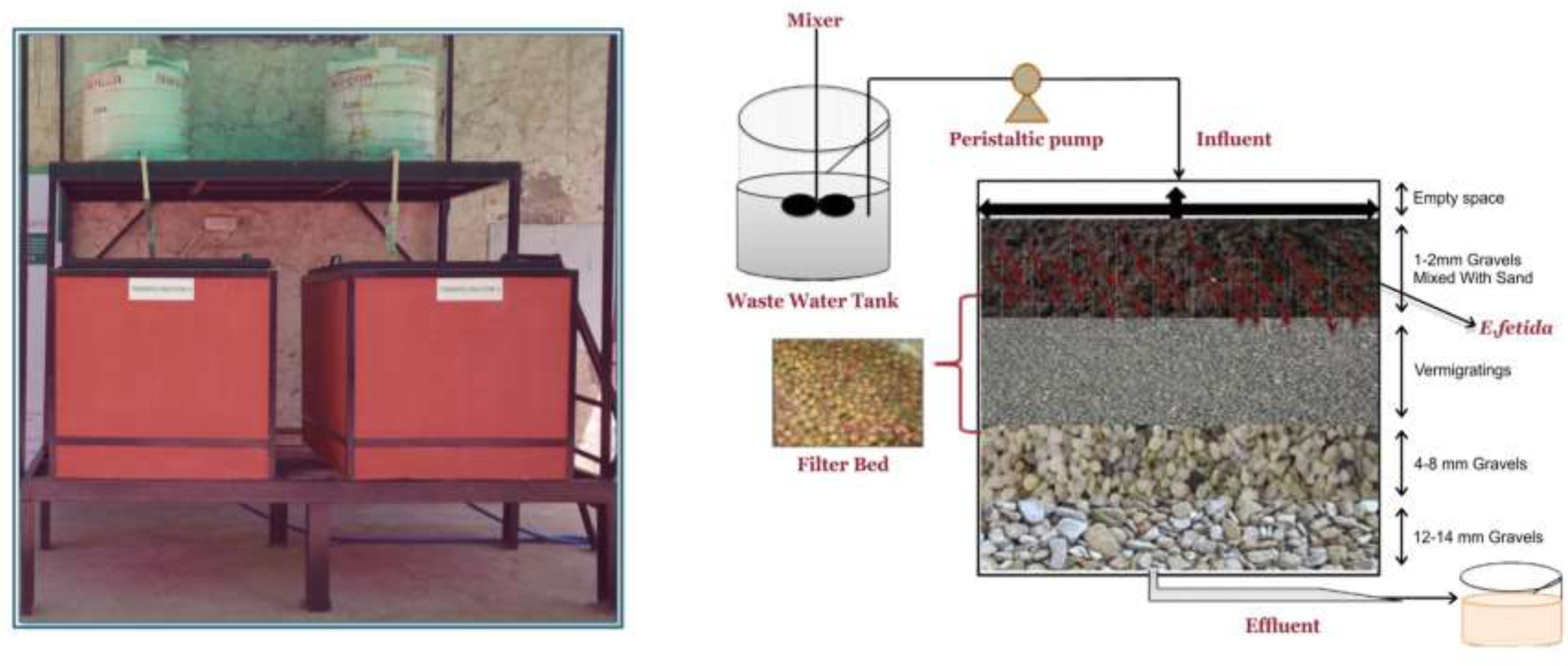
Field scale VF installed at the campus of Dr. B. Lal Institute of Biotechnology and Schematic flow diagram with design parameters.

### 2.2. Physical–chemical analysis and sampling

A continuous sampling was done once a week to evaluate the performance efficiency of the VF. The influent and effluent samples were analyzed for various physical, chemical and microbiological parameters. Physico-chemical parameters such as pH, Temperature, EC, TDS, and DO were monitored with the help of Hanna multi-parameter kit (APHA, AWWA, WEF. 2012). TSS was measured as defined by Arora et al., 2016. BOD was measured by azide modification method and COD was measured through closed reflux spectroscopic method (Hach-DR 5000) (Arora et al., 2014a). Nutrients (NO_2_^-^, NO_3_^-^ PO_4_^-^) were quantified and analyzed according to the methods described by Arora et al., 2016. Microbiological parameters such as Total and Faecal Coliforms and Faecal Streptococci (TC, FC and FS) were also conducted by most probable number (MPN) method (APHA, AWWA, WEF. 2012). The removal efficacy (%) and log removal were calculated as described in Arora et al., 2014a.

### 2.3. Investigation of microbial community diversity

At the end of the experimental period, different samples such as influent (IF), VF treated effluent (EF), VF-filter media (active layer) (VFM), earthworm casts produced by the earthworms in the VF (E_cast_), and earthworms coelomic fluid (E_CF_) were collected to investigate the bacterial community diversity and analyse the effect of earthworms on wastewater treatment.

#### 2.3.1. Earthworm extracts preparation

Microbial community diversity from the earthworms, were investigated from three different sets: control earthworms (E_C_) (that were not added to the VF), EW after 180 days of inoculation inside VF (E_180_) and EWs after 360 days of inoculation (E_360)._ Around 15 healthy earthworms were selected at random from the VF to determine a relative value based on 24 h of operation. Earthworm casts were sampled as described by Zhao et al., 2010. E _cast_ was expressed in mg of dry weight of the cast per gram of fresh weight of earthworm per day. The earthworms were kept in a culture dish in the dark for 24 h to empty their gut content, presuming that the residence time in the gut of the earthworms is 24 h. Their excrement in the culture dish was then rinsed with distilled water and dried at 105°C for 2 h, after which the dry weight was measured. In addition, the sampled earthworms were washed and dried with paper towels and then weighed. Their excrement in the culture dishes and adhering to the earthworms was collected and evaluated for microbial community structure.

#### 2.3.2. Coelomic fluid extraction

Coelomic fluid was extracted from the three sets of earthworms (E_C_, E_180_, E_360_). Cold shock method was used because it is safe for the earthworm and the yield is high (Patil and Biradar, 2017). After extraction, all the earthworms were viable and the coelomic fluid was processed further for investigating microbial community diversity.

#### 2.3.3. Filter media, influent and effluent sample analysis

Filter media sample was collected from the top layer of VF, in a sterile autoclaved petriplate to isolate the earthworm-associated microbial community present in the VF as described by Arora et al., 2014a. Spread plate dilution technique was used to isolate microorganisms from the samples. Well-isolated colonies were selected and sub cultured to a new petriplate. Cells from new colony were then picked up with an inoculating needle and transferred to an agar slant for maintenance of pure culture. Pure bacterial culture were identified morphologically (shape, gram staining and motility) and biochemically (sugar utilization, indole production, citrate utilization, methyl red-voges-proskauer test (MR-VP), triple sugar iron (TSI) utilization, oxidase production, catalase production, coagulase test according to *Bergey’s Manual® Syst. Bacteriol*., 2010. These were also further confirmed by Vitek identification tests.

### 2.4. Determination of Antimicrobial Activity & Enzymatic activity

The isolated microorganisms were tested for antibacterial activity against the known bacterial culture by agar well diffusion method (Parekh and Chanda, 2007; Sethi et al., 2013). Twenty-four hours fresh cultures of gram-positive S. aureus (ATCC 29213) and gram-negative E. coli (ATCC 25922) were swabbed by sterilized cotton swab and lawns were prepared over the agar surface. The molten Mueller–Hinton agar (Hi-media) was inoculated with 100 μl of the inoculum (1×10^8^ CFU/mL) and poured into the petriplate. About 50 μl cell-free supernatant was added in the well and plates were incubated at 37 °C for 24h. Antibiotic streptomycin (50 mcg) was used as positive control for the experiment. After 24h, the zones of inhibition were measured in millimeter. Diameters between 12 and 16 mm were considered to be moderately active and with more than 16 mm were considered to be highly active. The experiment was done in triplicate and the mean values were recorded. Enzymatic activity was determined according to the following methods. Protease activity was determined according to sigma’s non-specific protease activity assay (LOWRY et al., 1951). Cellulase activity was determined according to the method described by (Ghose, 1987). Amylase activity was determined according to the dinitro salicylate method (Miller, 1959). All the samples were analyzed in triplicate and the results were averaged. Optical density (OD) of each sample with reaction mixture was taken in a spectrophotometer. Enzyme activity was expressed in units/mL.

### 2.6. Protein estimation

Protein was extracted by using TCA method and the estimation was done using was done by using the Lowery’s Method

### 2.7. Protein Profiling of Coelomic Fluid

Coelomic fluid was extracted by the cold shock method and protein was precipitated using ammonium sulfate. Total protein extracted was quantified spectrophotometrically by checking absorbance at 260 and 280 nm. Samples from coelomic fluid of earthworms from different time points of the “microbial-earthworm interaction” establishment were normalized before performing SDS-PAGE. Standard Laemmli system as stated in (Laemmli, 2011) was followed for SDS-PAGE analysis.

## 3. Results & Discussions

### 3.1. Performance efficacy of wastewater treatment by the VF

The VF operated steadily without clogging during the experimental period for one year, indicating that there was a dynamic equilibrium between the earthworms and microorganisms and the quality of treated effluent is high. This is confirmed by analysis of the influent & effluent quality from the VF. The results of the performance efficacy of the VF are described in Table 2. The pH of the influent ranges from 8.0-8.5. The pH of VF effluent increased initially during the treatment, then decreasing slightly, reaching out to be in neutral range signifying the natural inherent ability of earthworms to act as buffering agent and neutralizing pH. Temperature is one of the key factors that affect the efficacy of wastewater treatment by vermifiltration. Throughout the study period, the average temperature of the influent and effluent from VF was in the range of 25-30 °C. Gravel and sand filter media have a buffer capacity to withstand the variations of temperature (Sinha et al., 2008). It provides better resistance to the adverse impact of vermifiltration under lower and higher temperature conditions (Li et al., 2009). Consequently, it proves to be a suitable dwelling habitat for earthworm *E. fetida* to thrive and perform its function proficiently in the optimum temperature range (Tripathi and Bhardwaj, 2004). The DO is an important factor that signifies the environmental conditions prevailing inside the reactor and it is one of the significant parameters in outflow, because low oxygen water is toxic to aquatic organisms. The value of DO in the influent was in the range of 0-0.35 mg/L. In VF effluent, the DO was observed to be 3-5 mg/L. This suggests that earthworms are responsible for creating aerobic conditions inside VF by their burrowing action. The design of VF is such that oxygen could penetrate to the bottom, thus increasing the efficiency of treatment. Higher DO in VF effluent reduces the septic condition and brings a good chance of wastewater reuse for irrigation purposes (Holenda et al., 2008). The moisture content of filter bed used in vermifiltration is an important parameter influencing the growth of the surface-feeding (epigeic) earthworm species *E. fetida* (Savigny or Lumbricidae) since the earthworm’s body contains about 80% water (Gunadi and Edwards, 2003). The moisture content in VF during the study period was observed to be 85–90% throughout, which is an optimum range for earthworm growth and activity. In the influent the average EC was observed to 2500-2800 μS/m whereas it increased in the effluent to 3000-3050 μS/m. The increase in EC during the process might have been due to the loss of weight of organic matter and release of different mineral salts in available forms (such as phosphate, ammonium and potassium). Earthworms ingest solid particles of wastewater and excrete them as finer particles. These finer particles trapped in the voids of VF and caused high removal efficiency of TDS from wastewater. TDS was found to be decreased significantly in the effluent (50-100 mg/L) as compared to the influent (1000-1500 mg/L). During the treatment process, TSS concentration in VF effluent was reduced significantly by 90%. This may be attributed to the ingestion of organic and inorganic solid particles in wastewater by earthworms, which excrete them as finer particles (Arora and Kazmi, 2015).

**Table 1:** Description of Filter bed layers.

| **Layers** | **Filter Media** | **Particle Size** | **Depth** |
| --- | --- | --- | --- |
| *Empty Space* |  |  | 15 cm |
| Active Layer | | 600-800 $\mu\text{m}$ | 30 cm |
| Second Layer | Small riverbed gravels | 2-4 mm | 15 cm |
| Third Layer | Riverbed gravels | 6-8 mm | 15 cm |
| Supporting Layer | Large riverbed gravels | 12-14 mm | 25 cm |
| Total Depth |  |  | 100 cm |

**Table 2:**
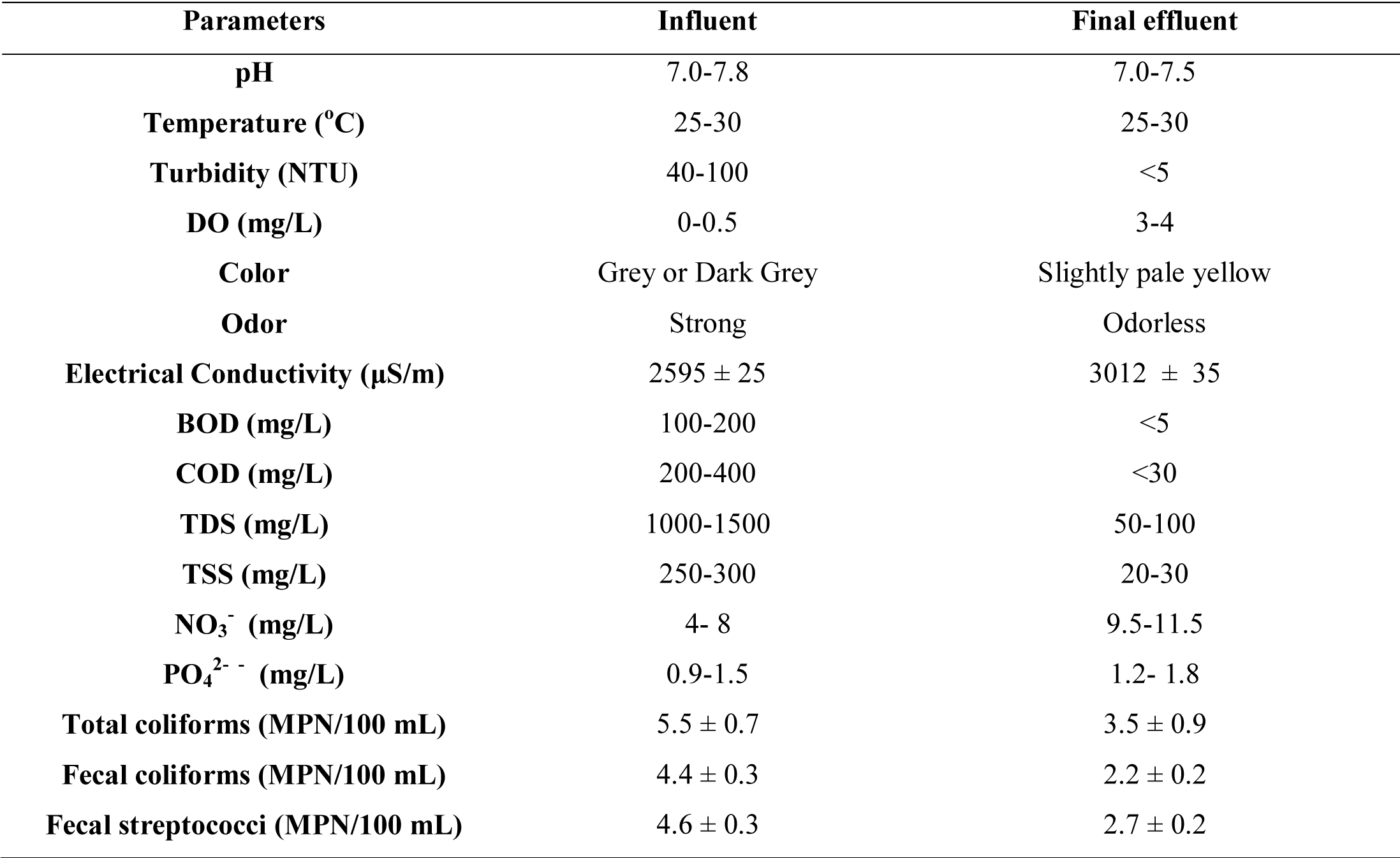
Physicochemical and microbiological parameters of influent and effluent samples.

| Parameters | Influent | Final effluent |
| --- | --- | --- |
| **pH** | 7.0-7.8 | 7.0-7.5 |
| **Temperature (°C)** | 25-30 | 25-30 |
| **Turbidity (NTU)** | 40-100 | <5 |
| **DO (mg/L)** | 0-0.5 | 3-4 |
| **Color** | Grey or Dark Grey | Slightly pale yellow |
| **Odor** | Strong | Odorless |
| **Electrical Conductivity (μS/m)** | 2595 ± 25 | 3012 ± 35 |
| **BOD (mg/L)** | 100-200 | <5 |
| **COD (mg/L)** | 200-400 | <30 |
| **TDS (mg/L)** | 1000-1500 | 50-100 |
| **TSS (mg/L)** | 250-300 | 20-30 |
| **NO<sub>3</sub><sup>-</sup> (mg/L)** | 4- 8 | 9.5-11.5 |
| **PO<sub>4</sub><sup>2-</sup> (mg/L)** | 0.9-1.5 | 1.2- 1.8 |
| **Total coliforms (MPN/100 mL)** | 5.5 ± 0.7 | 3.5 ± 0.9 |
| **Fecal coliforms (MPN/100 mL)** | 4.4 ± 0.3 | 2.2 ± 0.2 |
| **Fecal streptococci (MPN/100 mL)** | 4.6 ± 0.3 | 2.7 ± 0.2 |

These finer particles are further trapped in the voids of VF and cause high removal efficiency of TSS from wastewater (Sinha et al., 2008). This is the reason why VF do not choke but rather work smoothly and uninterruptedly.

The profile of BOD and COD for influent and effluent during the operation period shows that the organic matter measured as average BOD in the influent was 180-250 mg/L. The results show that earthworms can remove BOD loads by over 98% in VF to values less than 5mg/L. Higher BOD removal in VF is attributed to the activity of earthworms and its associated microorganisms that degrade the wastewater organics by its enzymatic activity (Rajpal et al., 2014; Sinha et al., 2008). The values of COD in the influent range from 325-400 and the average COD removal in VF is over 92% to values less than 30 mg/L in the effluent. This is attributed to the enzymatic activity of the microorganisms, due to the presence of earthworms that helps in the degradation of organic chemicals that otherwise cannot be degraded by microorganisms alone. Collectively, these results indicate that microbes are responsible for the biochemical degradation of organic matter, while earthworms are important drivers of the process so far, as they enhance the biodegradation through their feeding, burrowing and casting behavior. Thus, adding earthworms to the filter bed does seem to enhance organic matter removal (Liu et al., 2012). These changes in the wastewater after treatment indicate that oxidation, dehydrogenation and stabilization of organic matter, and the transformation of unsaturated structures to saturated structures (Romero et al., 2007) occur during the degradation of organic matter.

Nitrogen is one of the predominant contaminants in sewage, and nitrates and ammonia result in water pollution and eutrophication (Liu et al., 2012). The average concentration of NH_3_^+^− N in influent was 4.5 mg/L. Wang et al., 2011 had investigated that majority of ammonical nitrogen NH_3_^+^− N was removed mainly in the first two layers of VF through rapid adsorption by biomass and filters, and adsorbed NH_3_^+^− N was subsequently converted to NO_3_^−^ -N via biological nitrification, which was carried out by aerobic autotrophic bacteria using molecular oxygen as an electron acceptor. Meanwhile, high surface DO concentration was beneficial for aerobic microbial survival, which in turn was advantageous for nitrification. In addition, NO_2_ − N concentrations remained low in effluent because of the role of nitrites as intermediates in nitrification (Wang et al., 2011). Nitrification coupled with denitrification appears to be the major nitrogen removal process involved in VF, whereas insufficient available organic carbon is considered to be responsible for inhibition of denitrification. The total phosphate (TP) concentration in effluent increased significantly in VF. The change in influent and effluent TP concentration during the experimental period depicts that during the vermifiltration process, the increased TP concentration was attributed to enzymatic and microbial action of earthworms. The earthworm activities and associated microbes in the VF bed promote rapid phosphate mineralization in the system, causing increased concentration of TP in VF effluent (Kumar et al., 2015). Another reason behind the increased concentration of phosphorus may be attributed to the leaching of vermicast (excreta of earthworms, which are a strong source of nutrients i.e. 1.16% nitrogen, 1.22% phosphorus, and 1.00% potassium, and mainly responsible for conversion of soil/organic matter into vermicompost) from the filter material to effluent of VF (Kumar et al., 2015). The VF process mineralizes the nitrogen and phosphorous in the sewage to make it bioavailable to plants as nutrients (Kumar et al., 2015; Sinha et al., 2008) and thus indicates the potential of the vermifiltration system for treatment of wastewater to be reused for irrigation. Earthworms also host millions of decomposer microbes in their gut and excrete them along with nutrients nitrogen and phosphorus in their excreta. This must be due to the feeding action of earthworms that helps to control the over accumulation of biomass in the VF since earthworm burrowing and casting activities were observed. These results suggested that earthworms had significant effect on the enhancement of wastewater treatment performance during vermifiltration. Additionally, nitrogen and sulfur are essential for growth of all organisms. Microorganisms in the filter media convert a part of the nitrogen and sulfur in excess sludge into new cellular material for energy and growth (assimilation), and convert another part of them into simple inorganic compounds such as nitrate, sulfate, etc. (dissimilation) to be discharged in the filter effluent. Therefore, the comparable change suggests that microorganisms are primarily responsible for the turnover of organic matter in the VF

Presence of coliforms is often used as an indicator of overall pathogenicity of the sample. Samples were also analyzed for indicator organisms, i.e., total coliforms (TC), fecal coliforms (FC) and Fecal Streptococci (FS) for determining pathogen removal efficiency during the treatment. The average values of TC, FC and FS in the influent were in the range of log 7, log 3, log 5, respectively and it was observed that 2-4 log removal in the effluent. It could be due to better environment in VF for their proliferation than removal. Previous studies have shown the microbes that earthworms leave behind are beneficial to the VF because they compete with pathogenic organisms for the limited nutrients. The possible reason for pathogen removal is attributed to the fact that these pathogens get subjected to various toxic and antibiotic secretions from the earthworms and associated microflora (Sinha et al., 2008) .So, there is a possibility of antibacterial activity of the microorganisms that inhibits or prevents the growth of pathogens during the treatment. The concentration of pathogens (FC and FS) was reduced considerably and within the WHO standards for irrigation. So the investigation of the antibacterial activity of the isolated microbial species is crucial to understand the mechanism behind pathogen removal. Other attributed factors affecting pathogen removal in vermifiltration includes property of filter media to retain pathogens during filtration, i.e. bacterial adhesion, unsuitable physicochemical environment for pathogen survival and predation of these pathogens to regenerate bed for further adhesion

### 3.2. Effect of earthworms on microbial community structure

A total of twenty-nine bacterial species were isolated and characterized from influent and effluent samples. Gram staining and light microscopy characterized the bacterial isolates, which showed composition of 50% gram-negative and 50% gram-positive bacteria in influent. After the treatment of wastewater via VF, a radical change in microbial community diversity was observed in the effluent. Upon characterization 93.3% were grouped as gram-positive and 6.66% were grouped as gram-negative bacteria. These bacterial species were identified by biochemical characterization (as shown in Table 3) and further confirmed by Vitek 2 and were identified as *Bacillus, Micrococcus, Staphylococcus, Proteus, Corynebacterium* sp. which were common to influent and effluent samples, however, some pathogenic strains like *Enterobacter, E.coli and Psuedomonas* were present only in the influent, and were not detected in the final treated effluent. This important finding gave a clear indication that the pathogens are considerably removed during the vermifiltration process in the final treated effluent.

**Table 3:**
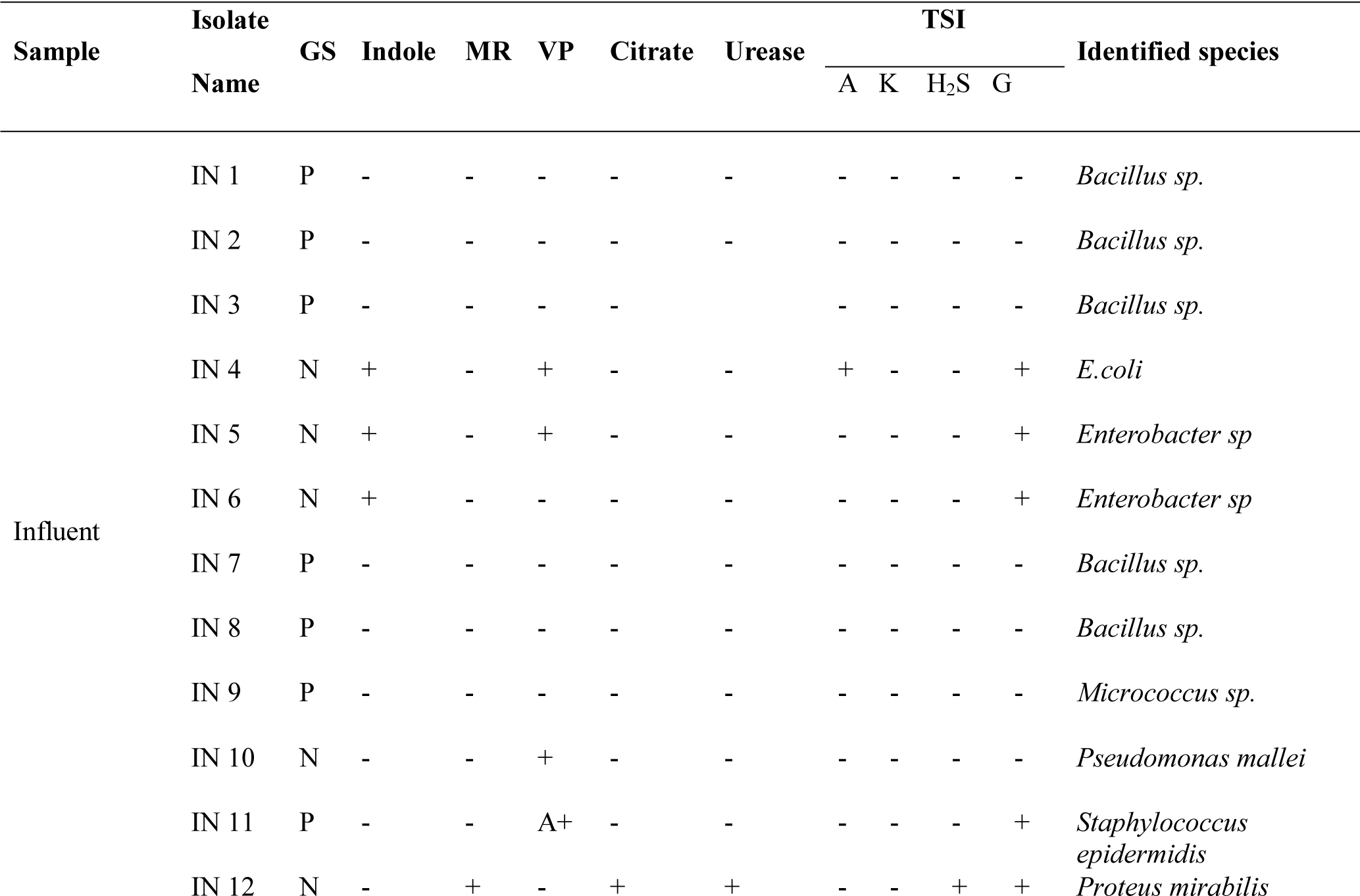
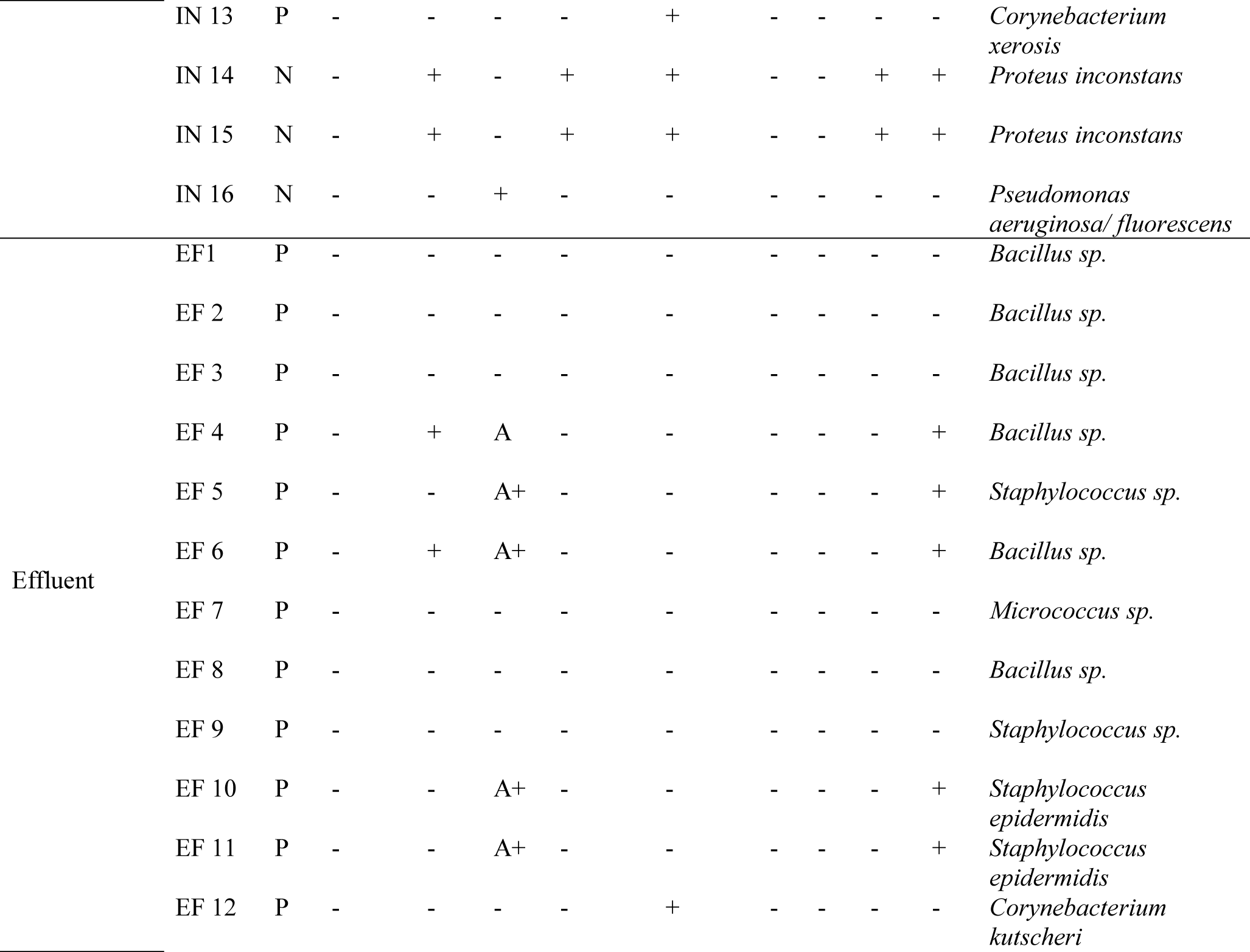
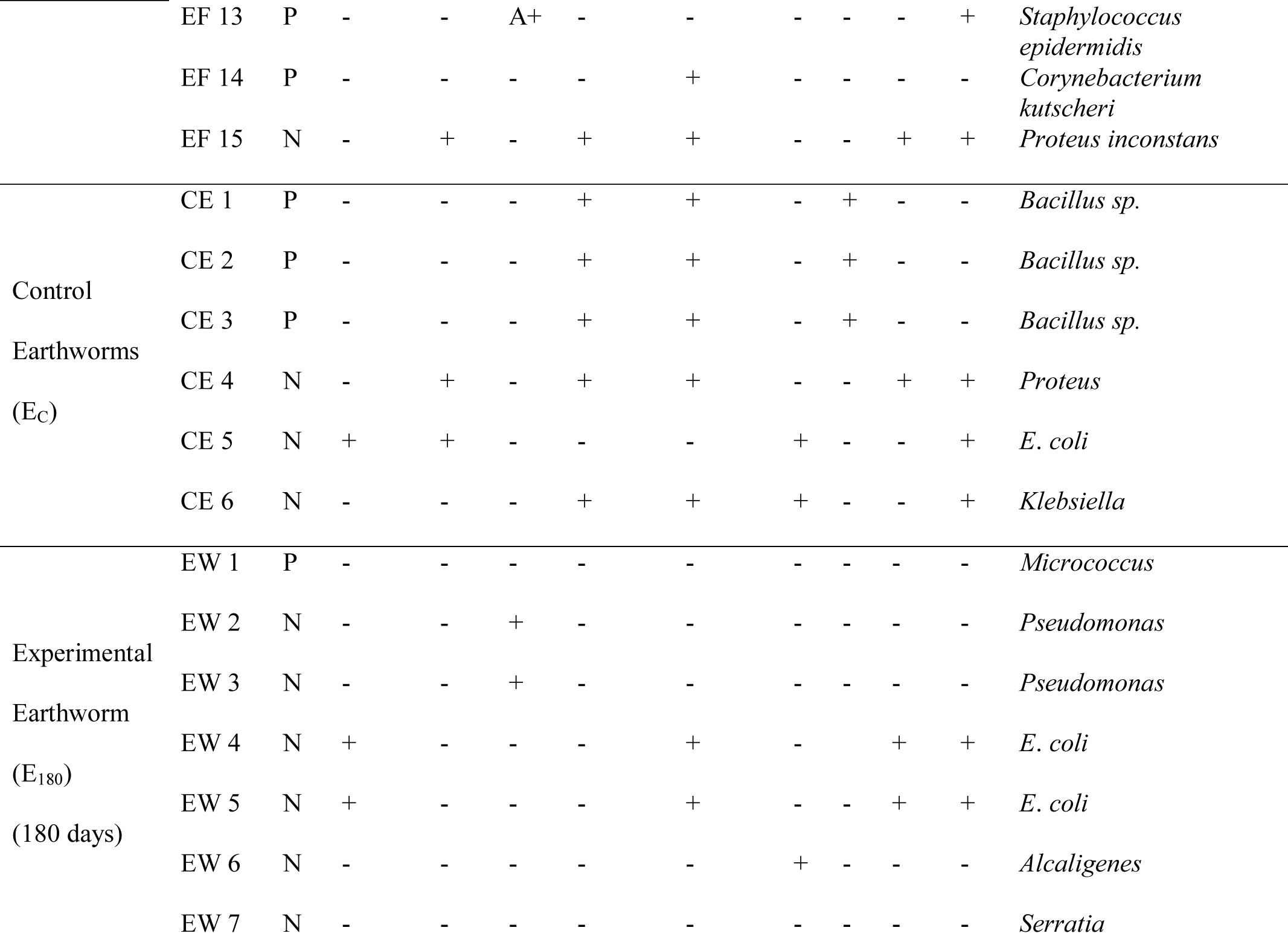
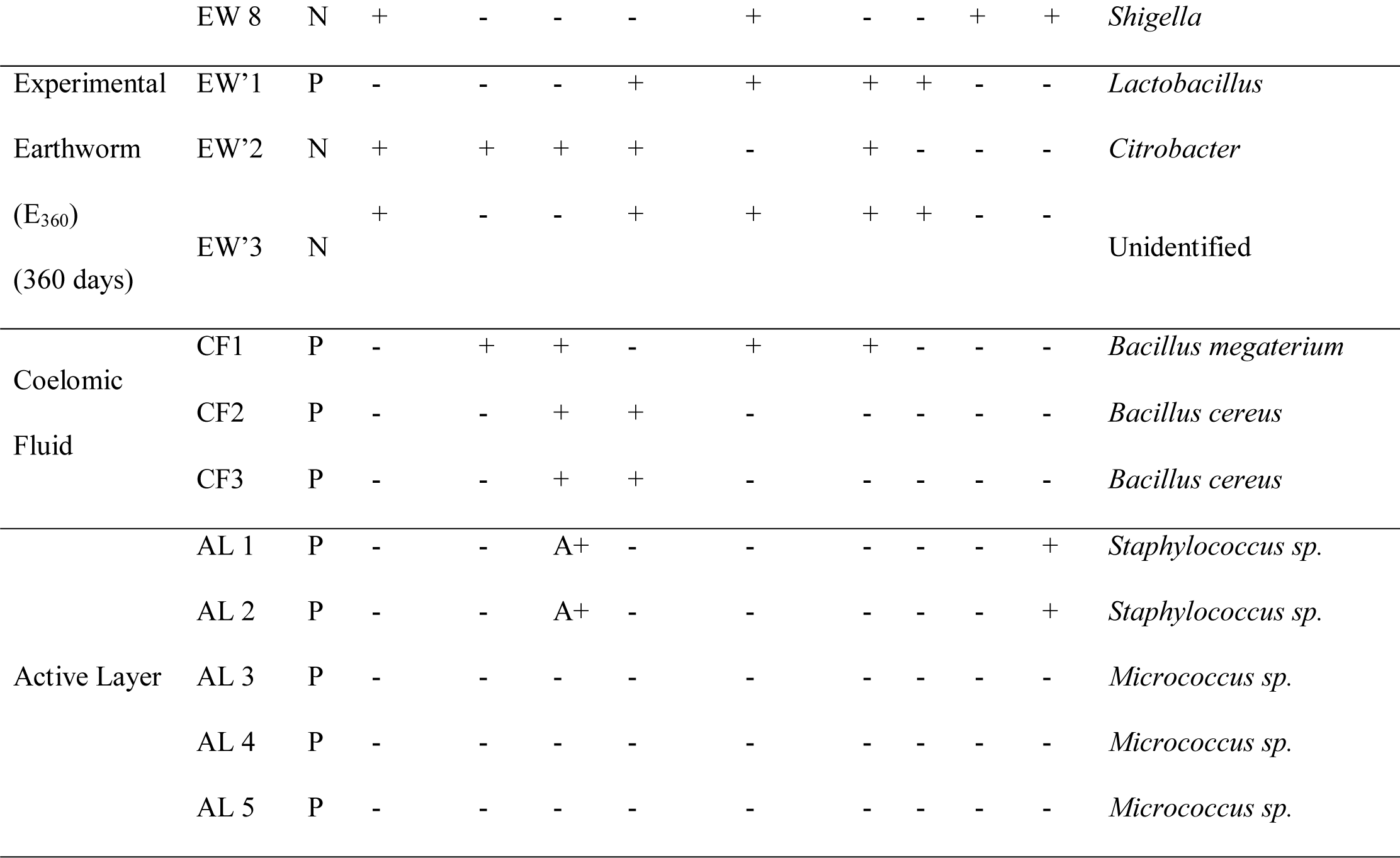
Identification and biochemical characterization of bacterial species from VF.

To further investigate about the obliteration of microbes inside the VF, earthworms and active layer samples were also investigated. Thirty bacterial isolates were detected from the earthworms and earthworm’s casts and active layers of the VF. Gram staining and light microscopy characterized the bacterial isolates, which showed the presence of 12.5% gram-positive and 87.5% gram-negative bacteria inside the earthworms (E_180_). The isolated strains were further depicted in Figure 2. Based on biochemical characterization, the dominant bacterial species identified were *Pseudomonas, E.coli, Micrococcus, Alcaligenes, Serratia* and *Shigella* in the earthworms. However, the results showed that in the filter media, the entire bacterial diversity showed presence of 100% gram-positive cocci, including species *Staphylococcus* and *Micrococcus* sp. Thus, these results implied that a part of Gammaproteobacteria might be the indigenous microbial species of earthworm gut which were consistent with the previous findings. Zhao et al., 2010 found that phylum Proteobacteria was an important contributor in vermifiltration. Li et al., 2009 reported that VF biofilms appeared to have more populations of Proteobacteria because indigenous gut-associated microflora from earthworms contributes to the microbial community inside VF. Studies have also shown that microbial community composition of earthworm gut changed with the growth substrate, implying that the earthworm gut microbial community had an interaction with that of the wastewater. Meanwhile, the changes in gut microbial community might affect the digestion of earthworms, and then caused an influence on the earthworm growth. There is a significant difference between the diversity of bacterial species present within the Earthworm (mostly Gram negative) and filter media active layer (Gram positive) thus, this supports the claim that EWs doesn’t only work as natural purifiers of the wastewater but also the soil within which they dwell. The top layer-vermicompost, can thus, safely be used as an agricultural byproduct in form of organic manure. This result is in consistent with the findings of Knapp et al., 2009, which suggested that an interaction between microorganisms and earthworms resulted in specific groups of bacteria being preferentially selected in the VF. By close analysis of the microbiota obtained from EW, we observed that they ranged from obligate aerobic (ex: *Micrococcus*) to facultative anaerobic bacteria (ex: *Serretia* and *Shigella*). This proves that even if during the whole treatment process, the reactor turns temporarily anaerobic, either because of clogging or disturbance in the regular HLR/HRT, the microbes present in the EW would excellently manage decomposition while the EWs restore the aerobic conditions through their burrowing activity. This result is in accordance with Arora et al., 2016, where they found the presence of an aerobic-anoxic zone in the VF. Certain *Pseudomonas* sp. are found to be plant growth-promoting and are used as biocontrols and bioremediation agents and *Klebsiella* reportedly fix nitrogen in the soil. Thus, their presence indicates enriched quality of compost formed as the result of the treatment. Some bacteria like *Citrobacter* use citrate as a sole carbon source. Their presence indicates the even decomposition of all the different types of organic matter. The findings indicate the existence of a special functional group of bacteria is suitable for the vermifiltration process.

**Figure 2:**
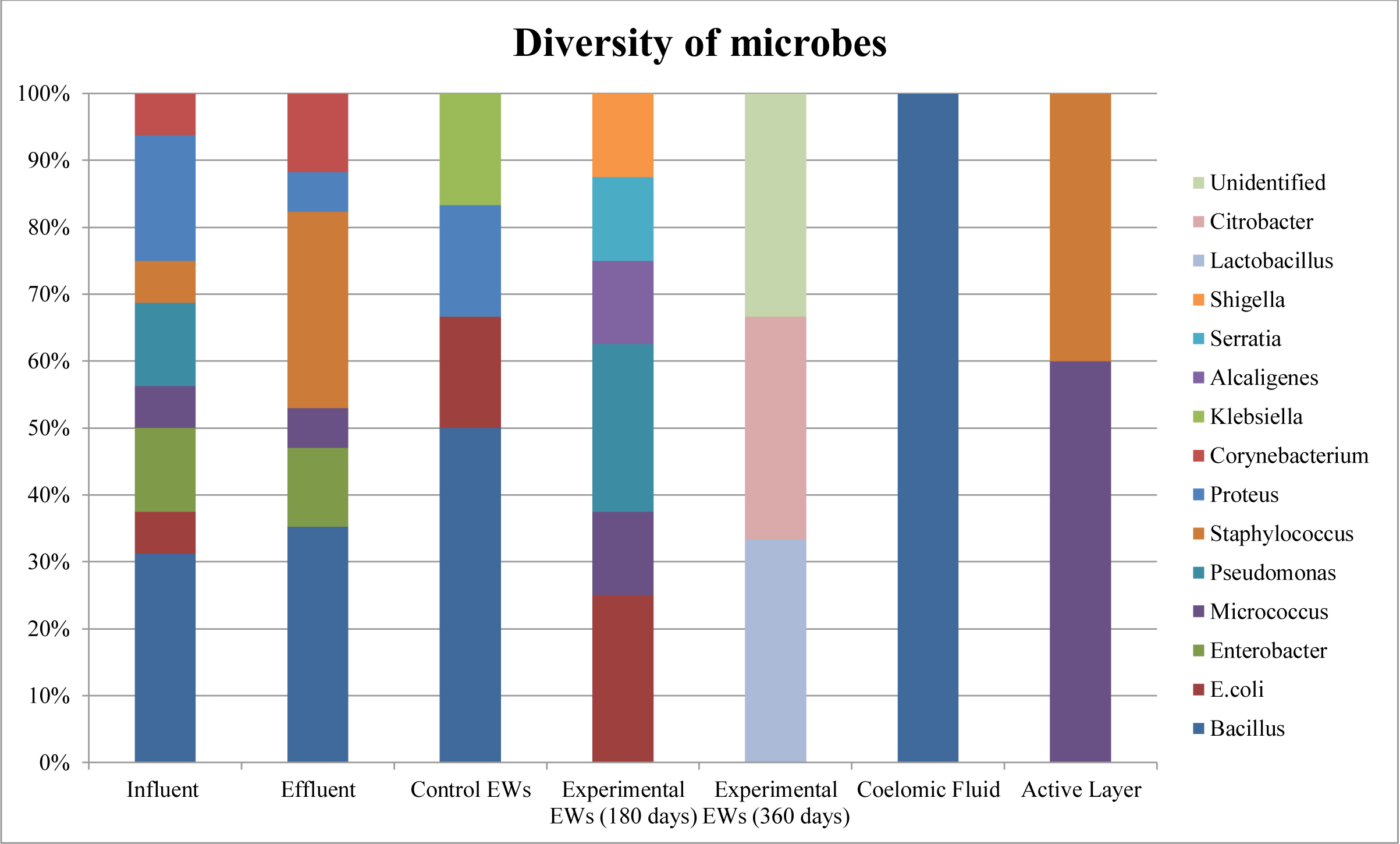
Percentage of Microbial community diversity isolated from various samples.

The results of the study further depicts the presence of three bacterial isolates isolated from the coelomic fluid, and were characterized to be Gram Positive *Bacillus* species. It is a notable contrast as the microbes isolated from EW gut are gram negative and in the CF and Bacillus appeared to be the dominant genus. These observations were consistent with the previous findings (Wang et al., 2011). This indicates that using earthworms in biofilters have a remarkable influence on the microbial community structure of the earthworms, filter media layer and the effluent, and that the vermifiltration process is an efficient technology for the wastewater treatment which is likely due to the presence of different microbial community inside the earthworms and active layer. These findings are in agreement with those of previous studies in which the passage of organic material through the earthworm gut resulted in stabilized earthworm casts (Aira and Domínguez, 2011). Specifically, earthworms in the VF were capable of transforming insoluble organic materials to a soluble form by the feeding action of earthworms, which led to the further degradation of organic matter by the microorganisms in the reactor. Collectively, these results indicate that microbes are responsible for the biochemical degradation of organic matter, while earthworms are important drivers of the processes in so far as they modify the microbial community structure through their feeding, burrowing and casting behavior. Thus, adding earthworms to the filter bed does seem to enhance excess wastewater treatment.

### 3.4. Effect of earthworms on microbial community function

The antimicrobial activity of isolated bacteria against known pathogens *S. aureus* and *E. coli*, is given in Figure 3. The antimicrobial activity was measured in terms of the zone of inhibition (in millimeters) around the tested organism. The control standard drugs gave an inhibition zone of 40 mm. It was observed that 69% of the isolates obtained from the influent, showed higher antimicrobial activity against both pathogens. These isolates were regarded as active strains while the percentage of active strains in the effluent decreased to 27%. Thus, it can be interpreted that the active microflora is present in the influent, that helps in the treatment process, particularly in removal of pathogens.

**Figure 3:**
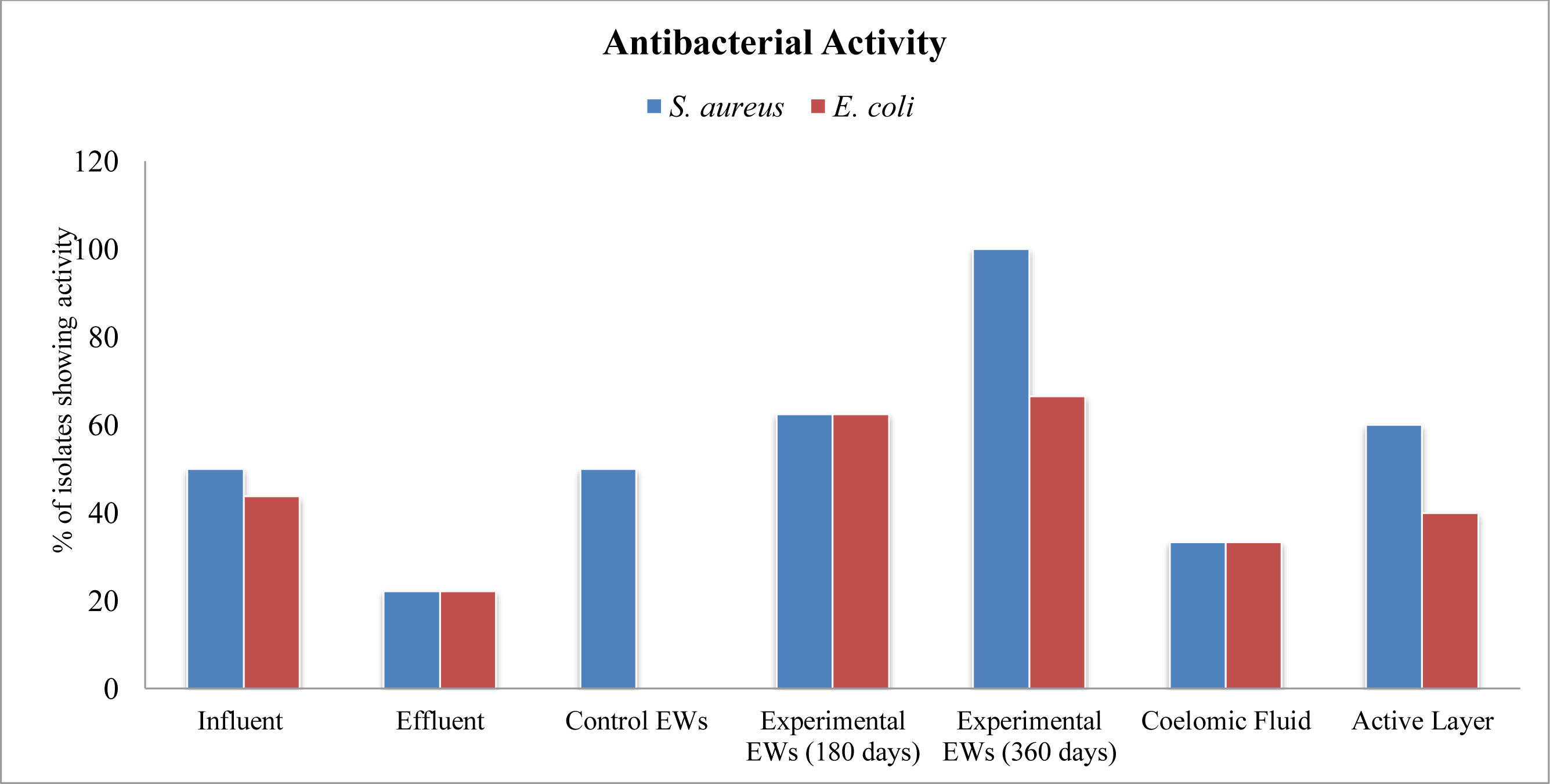
Percentage of bacterial isolates showing antibacterial activity against ATCC pathogens.

The results of the study also showed that 50 % of the isolates from E_C_ showed higher antimicrobial activity while in the case of E_180_ and E_360_ all the isolates (100%) showed the antibacterial activity. This clearly depicts that earthworms experience a significant change in their structure and function when exposed to the organic matter rich environment inside the VF, and they get adapted to the environment after their acclimatization period. The high antibacterial activity data of these active microbes can decipher the probable reason for removal of coliforms & pathogens from the wastewater. The filter media layer isolates also showed about 60% activity, which added a significant contribution to the pathogen removal.

The antibacterial activity of coelomic fluid showed variable results. It is interesting to observe that no antibacterial activity was seen at the concentration of 20μl but it showed slightly at concentration of 40μl in case of experimental EWs whereas in control EWs, it did not appear till the concentration of 60 μl. This again indicates that the EWs go through changes in their intrinsic microbial community or protein content as they adapt to the environment of the VF. The microbiota present in the coelomic fluid showed 33% antibacterial activity against the pathogenic strains. The probable reason is EWs prevails coelomocytes located in the coelomic fluid which is responsible for both innate cellular immune functions such as phagocytosis and encapsulation against pathogenic microorganisms and humoral components including lectin, preforming proteins, phenoloxidases and proteases nullifies antigenic material by agglutination, cytotoxicity etc. (Patil and Biradar, 2017). Thus, the enhanced activity of experimental earthworms signifies that a positive association is formed between EWs and the associating microflora of the active layer in which they dwell. This symbiotic association leads to biofilm formation which helps in the enhanced treatment of wastewater.

The production of enzymes by microorganisms which are found to degrade and stabilize the organics in wastewater is important due to their ability to decompose cellulose, proteins, starch and sugars, which guarantee the integrity of the vermifiltration system. The investigation on the enzymatic activity of the isolated bacterial species would supply vital data for understanding organic matter degradation in vermifiltration. The enzymatic activity of selected isolates is represented in Figure 4. Qualitative and quantitative assays were examined to detect the presence and absence and the amount of enzymes in the isolated microbial strains. These enzymes are responsible for the degradation of organic matter and decrease in the organic load of wastewater. A relatively high enzymatic activity is observed in the EWs associated microbes with all the isolates showing presence of any of the three enzymes. This finding also comes as an evidence for the presence of EW-microorganisms symbiotic relationship and is the probable reason for organic load removal during VF treatment. Earthworm–microorganism interaction promotes the organic matter degradation through improving the microbial activity and composition. Quantitative tests were performed for the isolates which were positive in the qualitative test to completely detect the magnitude of enzymatic activity. From the results, it was found that the enzymatic activity of the isolated microorganisms was very significant, resulting in organic matter biodegradation and the probable reason for higher BOD and COD removal

**Figure 4:**
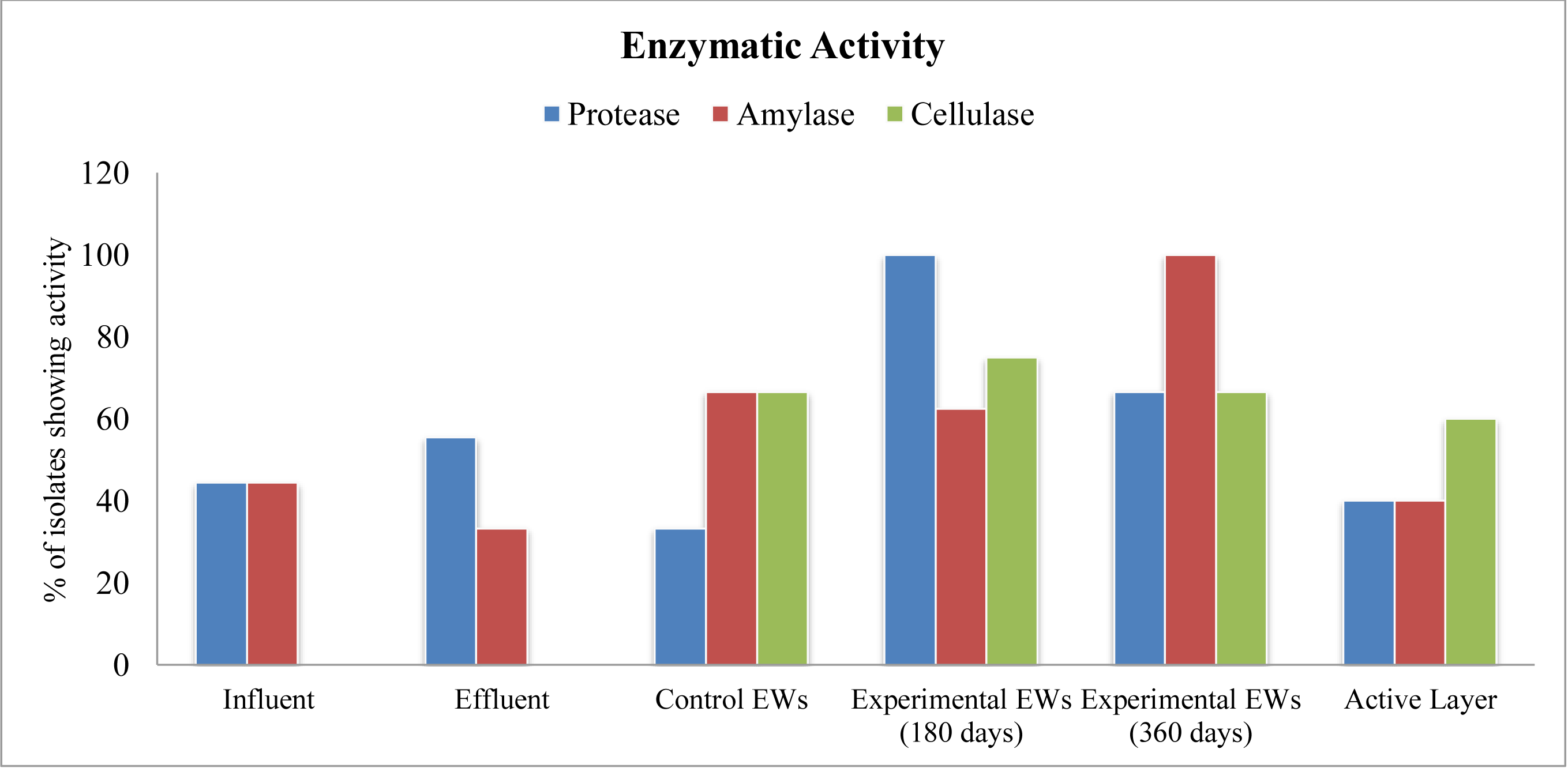
Percentage of bacterial isolates showing enzymatic activity.

### 3.5 Protein Profiling of Coelomic Fluid samples

The difference in the activity in coelomic fluid samples from different stages of earthworm microbial interaction establishment indicated that the EWs might be going through changes in the protein content of their coelomic fluid as they adapt to the environment of the VF. In order to confirm this hypothesis, protein was extracted from the coelomic fluid and protein expression pattern was analysed. Protein estimations showed that approx. 40.6 mg of total protein content was extracted from control EWs. However, in the case of experimental EWs only 31.9 mg of total protein content was observed. This data validates with the total yield of protein obtained. The data shows that, 2.70 mg/g and 1.772 mg/g of total yield was obtained from control EWs and experimental EWs, respectively. The changes in the habitat of the earthworms might be the possible reason for this change in protein content. Furthermore, the results of SDS-PAGE analysis showed the presence of distinguished bands for control and experimental EWs of molecular mass 48KDa, 35KDa and 25KDa. An additional band of 20KDa was also observed for control EWs, but was absent in the coelomic fluid of those experimental EWs which were under the process of interaction establishment. These results coincide with the previous studies of Rejnek et al., 1991; Tučková et al., 1991; Valembois et al., 1992 which demonstrated that chloragocytes present in the EW secrete two proteins with a molecular mass of 40KDa and 45 KDa, which share 35% similarities with immunoglobulins. It is also established that the chloragocytes synthesize 24 and secrete the protein lysenin of 33KDa, which binds specially to phospholipids of the cell membrane and causes cytolysis. Eiseniapore is also a 38-kDa haemolytic protein from *E. fetida* coelomic fluid that has been reported to bind to sphingomyelin and phospholipid vesicles indicating their role in lysis of membranes (Hatti and Ramkrishna., 2011). Thus, the presence of these protein bands observed in the coelomic fluid samples from the EWs established in VF study coincides with the previous findings and validates the role of microflora interaction in EWs physiology.

Combining the above results with the previous deduction that the treatment efficacy of VF is highly dependent on the antimicrobial and enzymatic activity of EWs, the role of earthworms in VF treatment plants seems to be significant. However, it is also clear from the experiments checking for anti-microbial and enzymatic activity that the isolates from the gut of established EWs were able to perform significantly better as compared to the isolates outside the EWs gut. Thus, it can be concluded that the earthworms and associated microbes both are acting in a combination as the factors responsible for the treatment of wastewater. Microorganisms play a very significant role in the pathogen removal mechanism and organic matter reduction in response to their interaction with the earthworms. This technology thus skillfully incorporates the individual potential of microbes and earthworms.

## 4. Discussions

Vermifiltration is a nature based sustainable technology which integrates the potential of earthworms and gut-associated microorganisms and microbes dwelling in the filter bed for the wastewater treatment process. The present field scale study was conducted to analyze the institutional wastewater treatment competence of a vermifilter, installed at the campus of Dr. B. Lal Institute of Biotechnology, Jaipur. The results of the study revealed significant removal efficacy of VF for BOD and COD and coliforms & pathogens. In addition to the treatment, this study also elucidates essential and effective microbiota responsible for the high removal rate via a thorough analysis of earthworms and filter media associated microbes, investigating their respective roles. To develop a comprehensive conclusion, we only considered the dominant strains and these results are illustrated in Figure 5. Based on the analyses, it was clear that earthworm activity had a profound effect on the physical and chemical composition of the treated wastewater, and that this effect occurred primarily via an increase in the organic matter available for the microbial degradation. The role of the the earthworms is clearly explained in the Figure 5. It is well known that microbial communities preferentially select soluble substances as their diet (Knapp et al., 2009). Earthworms are able to transform organic materials from insoluble forms to soluble forms that are available for further degradation by microorganisms, thereby enhancing the microbial population and activity. The passage of organic material through the earthworm gut resulted in physical decomposition due to the muscular grinding action of the gizzard (Lazcano et al., 2008). It has also been reported that the earthworm gizzard is a colloidal mill in which the feed is ground into particles. Additionally, the stimulatory effect of earthworms on the microorganisms could be explained by the mucus and casts that the earthworms produced. Mucus is a source of easily assimilable carbon for microorganisms, while casts are often enriched with available forms of C, N and P (Aira et al., 2007), and contain more active microbial communities than the foods that earthworms consume (Suthar and Singh, 2008). Furthermore, it has been established that *Eisenia foetida* has a unique indigenous gut-associated microflora that contributes to the development of a diverse microbial community in VF systems. The dynamics of EW and microorganisms for the possible removal of contaminants from wastewater is illustrated in Figure 6.

**Figure 5:**
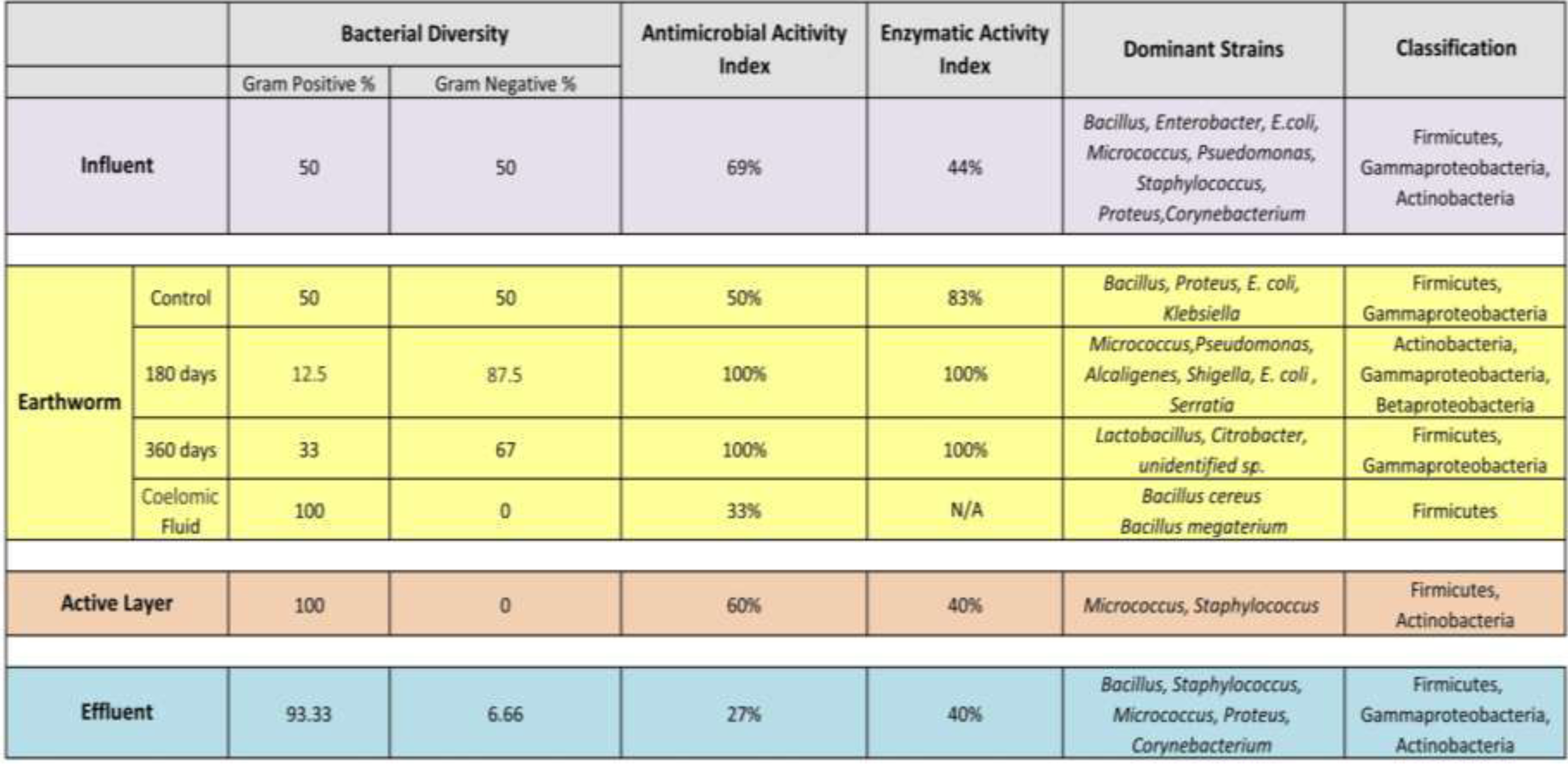
Percentage composition of microbial community diversity: structure & function in different samples.

**Figure 6:**
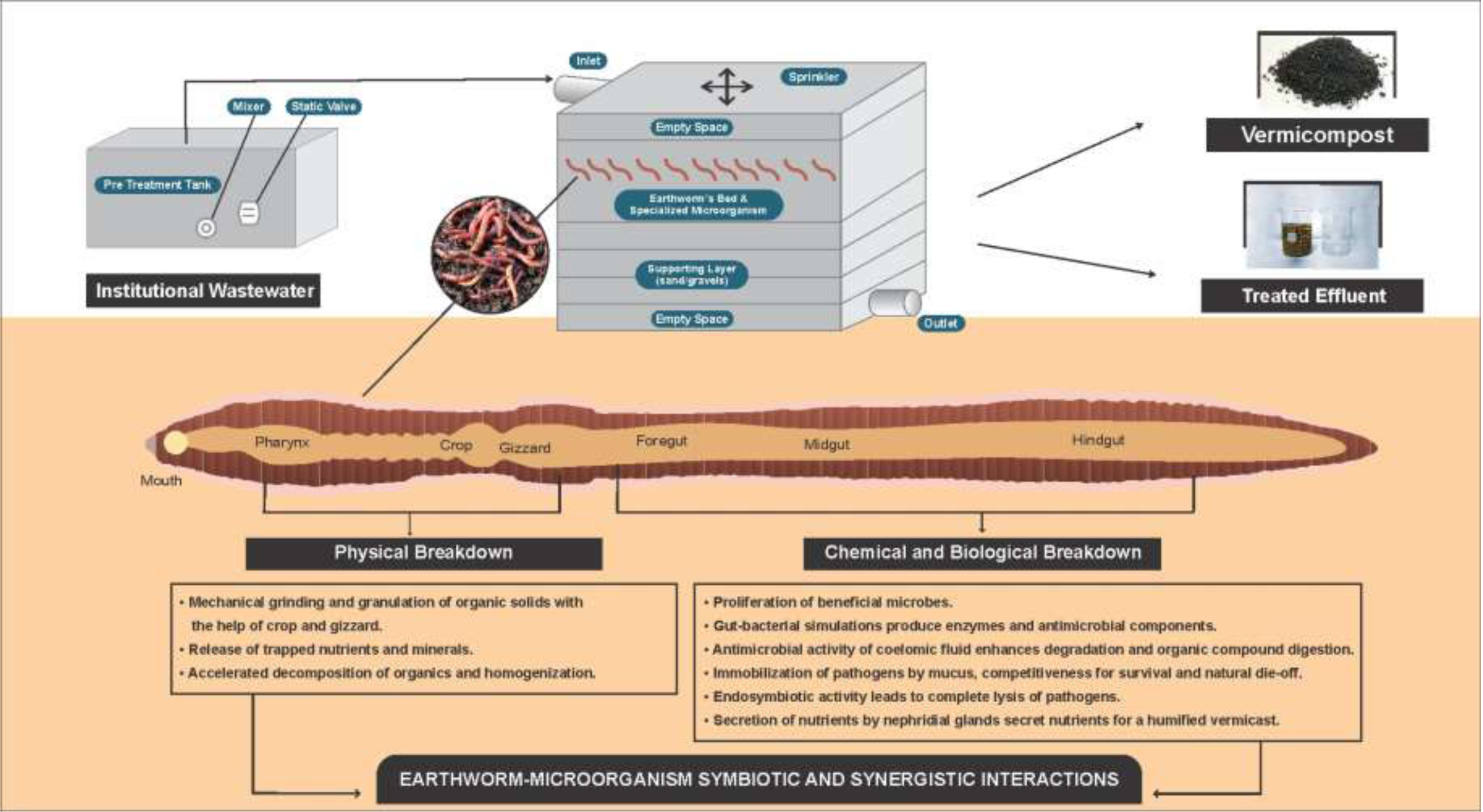
Adaptation of earthworms inside VF and mechanisms underlying treatment.

Previous studies have also revealed that during the symbiotic relationship of earthworms and microorganisms, microorganisms perform biochemical degradation of the waste material, while earthworms degrade and homogenize the material through muscular actions of their foregut and add mucus to the ingested material, thereby increasing the surface area for the microbial action (Aira et al., 2007; Suthar and Singh, 2008). A similar benefit was also obtained when earthworms were introduced into the VF systems in this study, suggesting a mutualistic relationship between earthworms and microorganisms. Thus, earthworm and microorganism interactions accelerated the mineralization of organic materials, favoring the breakdown of excreted polysaccharides. Taken together, vermifiltration process is predominantly driven by the earthworms and the earthworm– microorganism interactions played an important role and significant improvement in the wastewater treatment and stabilization.

There is a gradual increase in the structure of gram negative bacterial diversity of the earthworms when compared in control and experimental EWs (EW_control_ < EW_180days_ and EW_360days_). A similar trend can be seen for their functional aspects such as antibacterial and enzymatic activity. However, in active layer, the results were very astonishing as there was a complete absence of gram-negative bacteria diversity. There is a possibility that negative predation of EWs lead to high gram negative diversity inside the gut, cleansing the active layer and making it safe for agricultural use as manure. This has also been stated in previous studies that pathogens get devoured by earthworms and are killed in their gut. Mucus and CF have also been reported to support the survival of favorable microbes by selective ingestion and later eradication of pathogens in the alimentary canal. It produces antibiotic substances which do not favor the growth kinetics of the pathogens and immobilize them due to its sticky nature. Thus, the pathogens present in a hostile environment, halted and facing a lack of food, ultimately are destroyed off. It also helps in mineralization of the effluent and vermicast (Arora et al., 2014b; Singh et al., 2019, 2018). This can be clearly verified by the antibacterial activity of CF from experimental and control EWs (Table 4) as the EWs after treatment started responding to pathogens at lower concentration with high activity. However, it should also be implicit that earthworms alone are not capable of treating the wastewater with such efficiency. Microorganisms play a major role in pathogen removal efficacy in this technology as proven from the protein profiling assay.

**Table 4:**
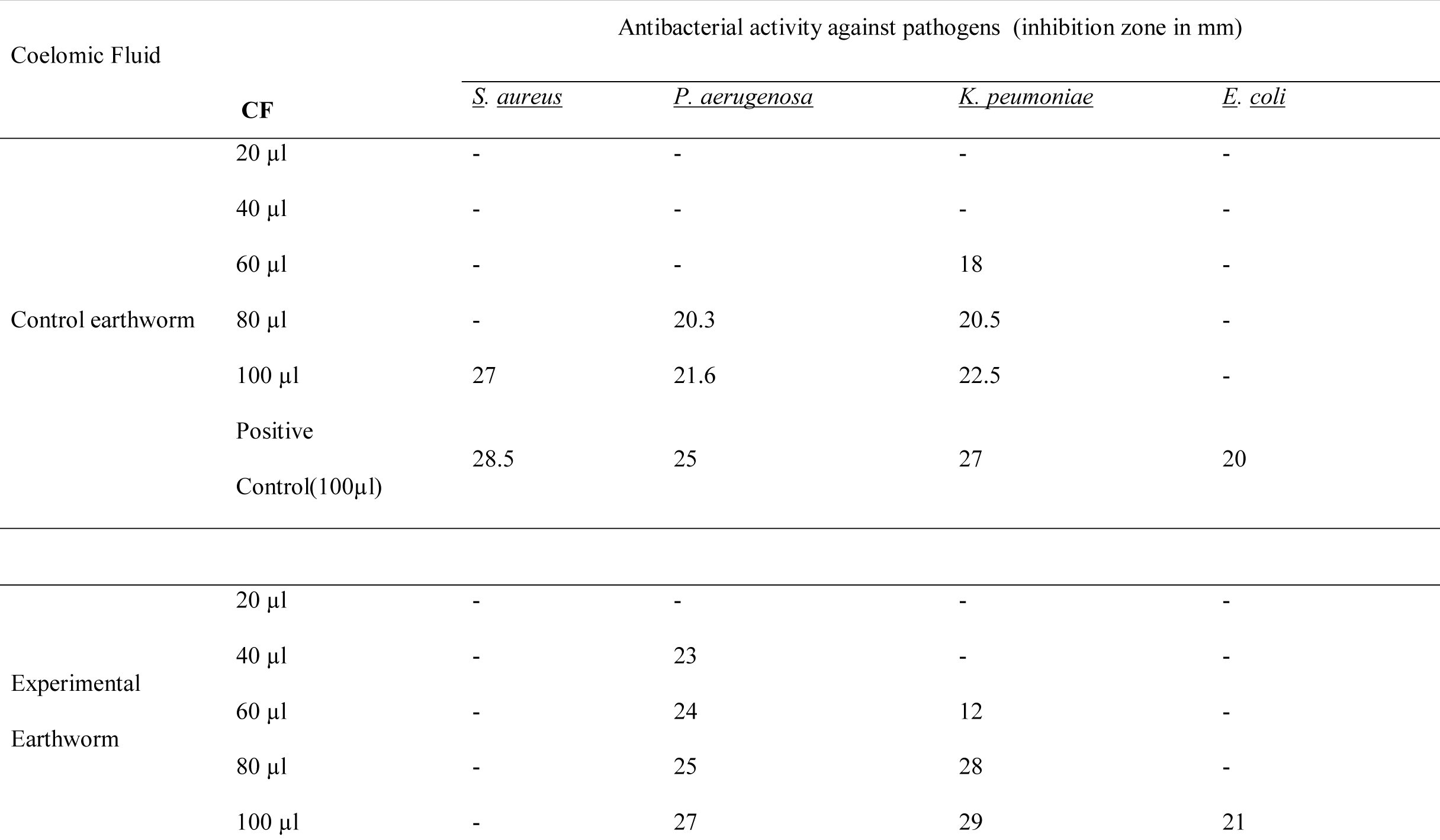
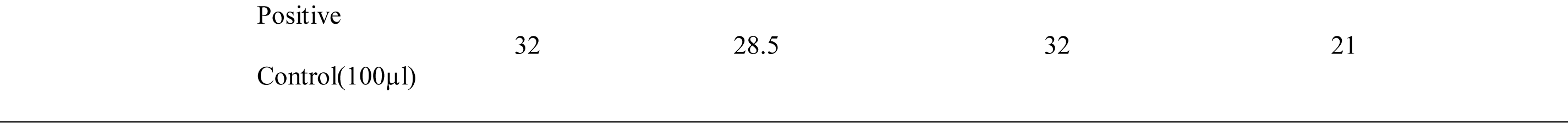
Antibacterial activity of coelomic fluid.

The microbes present in the EWs and filter media together lead to biofilm formation which acts upon the various components of wastewater such as TDSS, pathogens, organic matter and nutrients as depicted in Figure 7. Solids are broken down as a result of the catabolic activity. Organics are subjected to the enzymatic activity whereas pathogens are removed by the antimicrobial activity. Nutrients are recycled and added to the byproducts viz. vermicompost and vermiwash.

**Figure 7:**
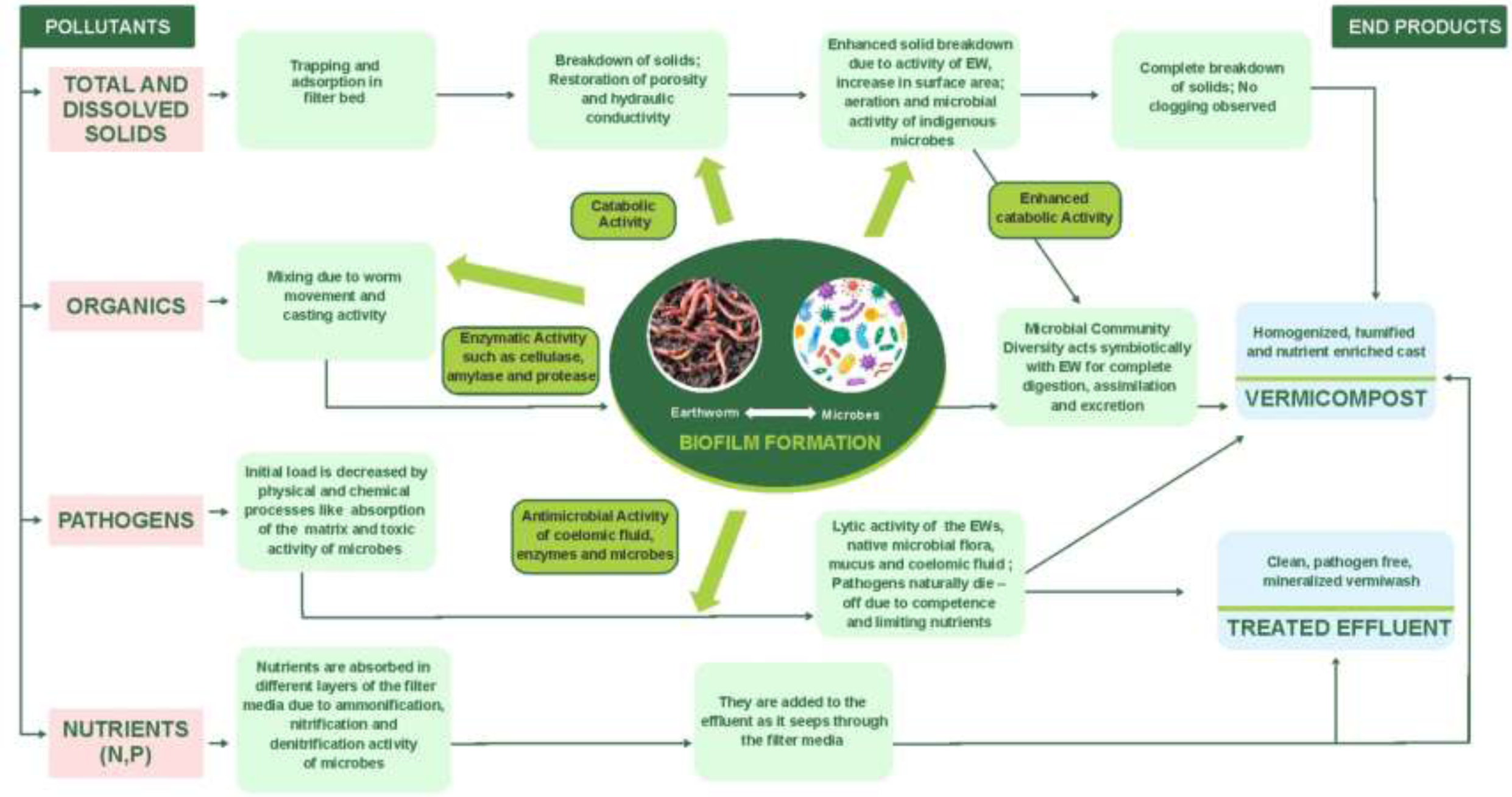
Earthworms-Microorganisms complex dynamics for wastewater treatment.

Thus, we can conclude that vermifiltration is an uncomplicated process when talked in terms of resources used to set it up, however, the whole organics and pathogen removal mechanism is quite complex as various dominant strains present at each level, contribute individually as well as in combination with the EWs coelomic fluid to helps in microbial load reduction and organic matter degradation.

## 5. Conclusions

The dynamics of earthworms and microorganisms interactions were studied for the first time, in a field scale vermifilter and established that earthworms had a significant effect on controlling microbial biomass and improving microbial activity. In addition, microbial community dynamics analysis for the influent, effluent, active layers, and earthworms demonstrated the bacterial community structure and functions to understand the correlations & symbiosis to decipher the mechanisms for wastewater treatment that played an important role earthworm regulation. Moreover, as earthworms emerge as important biological invaders, the results of the present study may help to fully appreciate, estimate and model the consequences of this momentous global change phenomenon. Particularly, the spread of beneficial microorganisms inside the VF, and earthworm species likely threatens the potential pathogens diversity and functions. Some bacterial groups resided preferentially on the VF biofilm, while earthworm cast played a potentially significant role in the carbon and nutrient removal processes. These results indicate that earthworms help to guarantee the efficiency and stability of VF performance for the treatment of wastewater. The resulting end products treated effluent and vermicompost can be used for sustainable agriculture. This study can become the foundation for further developing vermifilters as a potential nature based solutions.

## Credit authorship contribution statement

**Sudipti Arora**: Conceptualization, Investigation, Resources, Manuscript Writing and editing, corresponding author

**Sakshi Sarawat:** Experimental design, protocol standardization, Experimental conduction, Data analysis, Writing -original draft of manuscript, Investigation

**Rinki Mishra** Experimental design, protocol standardization, Experimental conduction, Data analysis

**Aditi Nag**: Experimental design on molecular work, protocol standardization, Experimental conduction

**Jasmine Sethi:** Sampling, Experimentation, Data analysis

**Jayana Rajvanshi**-Sampling, Experimentation, Data analysis

**Sonika Saxena:** Resources providing, Sample collection, and supervision

## Acknowledgements

The study group would like to acknowledge the constant support received from Dr. B. Lal Gupta (Director, Dr. B. Lal Institute of Biotechnology, Jaipur) and Dr. Aparna Datta for inspiring this research and providing daily motivations to work faster. The team further acknowledges the efforts made by research students Ms. Kirti Agarwal, Ms. Sonit Kumari, Ms. Anamika Verma and Ms. Shruti Kakkar, for conducting experiments.

---

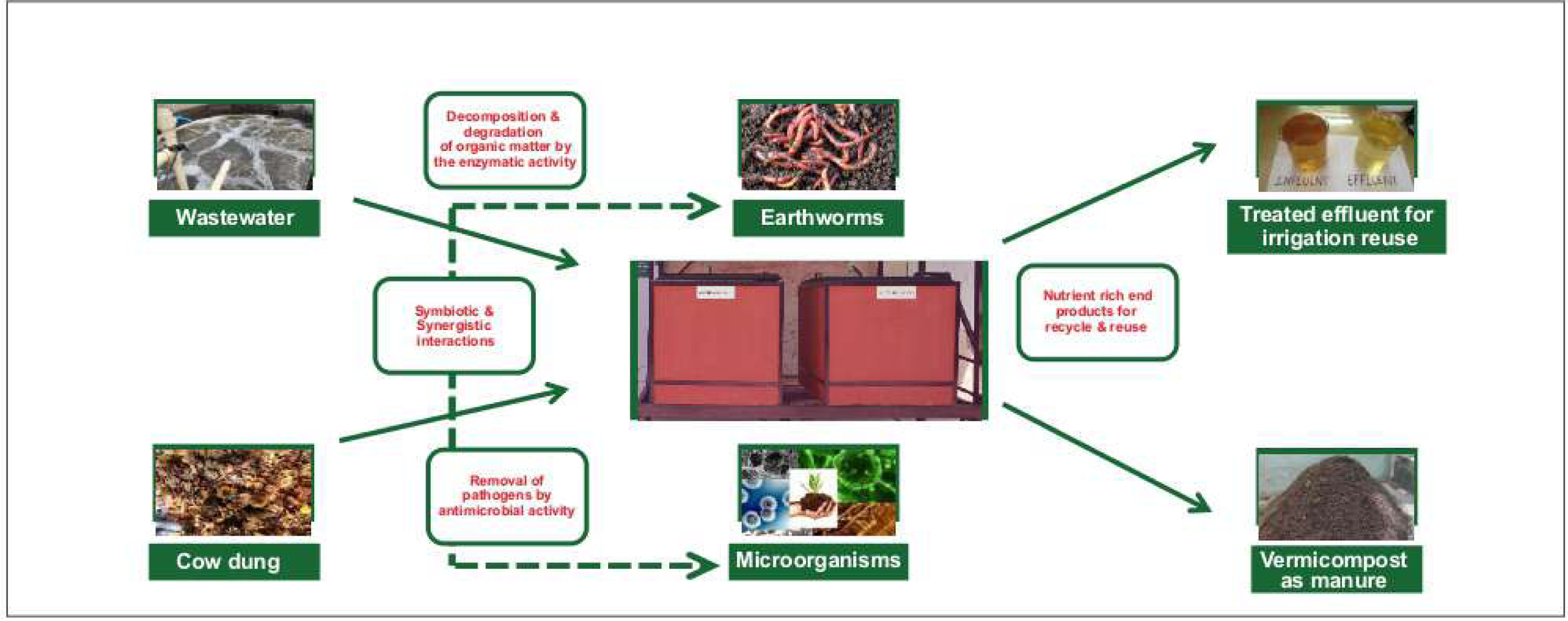

## Highlights

- Vermifilter is able to efficiently treat institutional wastewater.
- Earthworms-Microorganisms interactions are responsible for wastewater treatment.
- Investigation of the dynamic mechanisms of microbial community diversity inside a VF was explored.
- Earthworm gut microbial communities were dominated by *Gammaproteobacteria*.
- Filter media layer showed presence of *Firmicutes* and *Actinobacteria*.

